# Advancing Plant Health in Sub-Saharan Africa: Understanding National Agricultural Research Institutions capacities for Strategic Investment Priorities to Boost Genetic Gains Against Crop Biotic Stress

**DOI:** 10.64898/2026.07.19.739424

**Authors:** Janet Kimunye, Boubacar Sinare, Hermann Some, Inoussa Drabo, Tamerlane Mark Nas, Abhishek Rathore, Roma Das, Baloua Nebie, Haile Desmae, Joseph Bigirimana, Ajay Panchbhai, Harish Gandhi, Amos Emitati Alakonya

**Author notes:** **Correspondence:** Amos Alakonya A.

## Abstract

Plant health research within National Agricultural Research and Extension Systems (NARES) represents a critical yet neglected pillar of agricultural resilience in sub-Saharan Africa (SSA), where crop productivity remains substantially below global averages due to persistent biotic and abiotic stresses. Although plant health units have a strategic mandate to support crop protection, disease surveillance, and varietal improvement, the institutional capacity of these units across NARES in SSA is not defined. Here, we assessed 36 plant health units across 26 SSA countries to determine research capacity, identify operational constraints, and examine their contributions to crop improvement and food security. Our findings reveal the existence of a strong human capital operating under structurally constrained systems. More than 65% of personnel possess postgraduate qualifications, indicating the presence of a highly trained scientific workforce with considerable potential to drive plant health innovation. However, this expertise is undermined by severe infrastructural and institutional limitations that restrict research delivery. Access to essential facilities remains low, whereby only 53% of respondents had access to plant pathology laboratories, 43% had molecular biology platforms, and 37% had glass/screenhouses, while only 30% of respondents maintained phenotyping infrastructure such as sick plots or endemic/hot spot disease sites. These deficiencies undermine plant health research in areas like biotic stress resistance screening, pathogen diagnostics, and diversity. It was noted that plant health units primarily target staple food crops central to regional food systems, including cereals (maize, rice, sorghum, and millets), legumes (groundnut and cowpea), and root and tuber crops (cassava and potato). All these crops are affected by diverse biotic stresses that include fungi, bacteria, viruses, insect pests, and parasitic weeds. Yet research effectiveness to address these challenges is further constrained by unclear or fragmented protocols, poorly equipped laboratories, weak data capture, and management systems. Critical gaps were consistently identified in pathogen isolation and characterization, disease phenotyping, surveillance and mapping, experimental design, and data management. Workforce composition raises additional concerns regarding long-term system sustainability. Women account for only 18% of plant health personnel, while researchers younger than 35 years represent just 10% of the workforce, reflecting weak generational succession and gender inequity. These demographic imbalances constitute an existential threat to institutional continuity and innovation capacity. In summary, closing these gaps will require coordinated investment in infrastructure, technical training, institutional strengthening, and digital modernization. Equally important is the development of inclusive workforce strategies that improve gender representation and strengthen succession pathways. Unlocking the potential of NARES plant health networks is essential for accelerating resilient crop development, strengthening agricultural productivity, and advancing food security across SSA.

## Introduction

Sub-Saharan Africa (SSA) faces a critical challenge of sustainably increasing food production to meet the demands of a rapidly growing population. Despite its agricultural potential, the region continues to experience persistently low productivity relative to global averages, contributing to chronic food insecurity (Thomas, 2020). Large and persistent yield gaps between actual and attainable production underscore the need for transformative approaches to agricultural intensification (Ittersum et al., 2016; Tittonell & Giller, 2013). Among the key constraints, biotic stresses including fungi, bacteria, viruses, phytoplasmas, nematodes, insects, and parasitic weeds are the major cause of crop losses across farming systems in (Day et al., 2017; Yali, 2022). The magnitude of these losses is substantial; for instance, wheat rust epidemics can reduce yields by up to 70% (Singh et al., 2016), rice blast by as much as 50% (Nalley et al., 2016), and Fusarium wilt in pigeon pea can lead to complete crop failure (Choudhary et al., 2013). These challenges are further intensified by the emergence and spread of new pests and diseases, driven by globalization, climate change, and agricultural intensification (Pixley et al., 2023). As a result, the risk landscape for crop production is becoming increasingly complex, undermining gains from past agricultural investments. Addressing these challenges requires a more integrated approach that embeds plant health and phytopathology within crop breeding, agronomy, and pre- and post-harvest systems to enhance resilience and productivity.

Historically, efforts to improve agricultural productivity in SSA have largely emphasized the adoption of external inputs such as fertilizers and improved seeds (Thomas, 2020). While important, these approaches alone have not delivered sustained productivity gains. There is increasing recognition that long-term improvements depend on strengthening research systems capable of generating, adapting, and scaling context-specific agricultural innovations (Mokwunye, 2010). Conventional metrics of research performance, such as publication output (Bennell & Thorpe, 1987), are insufficient proxies for impact. Instead, there is a need to shift toward integrated frameworks that prioritize human capital development, infrastructure strengthening, and interdisciplinary collaboration across plant health, crop improvement, and agronomy. Such a transition is essential to generate scalable and sustainable solutions that directly address food security challenges in SSA.

Global frameworks such as those advanced by the International Plant Protection Convention (IPPC) provide guidance for pest risk analysis and plant health management (Carvajal-Yepes et al., 2022; Prasanna et al., 2022). However, the effective implementation of these frameworks in SSA is constrained by limited technical capacity, inadequate infrastructure, and weak institutional coordination. In many cases, plant health research efforts remain fragmented, with limited collaboration among national agricultural research and extension systems (NARES) and international partners, and a lack of harmonized technical and operational frameworks (Stads et al., 2022). This fragmentation has slowed the development, validation, and deployment of plant health innovations across the region.

To address this gap, we conducted a comprehensive assessment of plant health research capacity within NARES across SSA. The objectives of this study were: 1. To assess plant health research capacity across NARES in sub-Saharan Africa, focusing on human resources, infrastructure, and system functionality 2. To identify key constraints and investment priorities for strengthening biotic stress research and its integration into resilient, food-secure agricultural systems. Overall, this study provides a data-driven regional roadmap for strengthening plant health systems in sub-Saharan Africa, identifying priority investments in infrastructure, workforce development, digitalization, and coordination needed to enable effective biotic stress phenotyping that will accelerate genetic gains for food security.

## Materials and Methods

### Survey

An online survey was designed by a team of breeders and pathologists from CGIAR and NARES centers. The questionnaire (Supplementary 1) was administered to 57 crop protection contact scientists from 42 institutions across 26 countries (Supplementary 2). Survey questions were designed to understand the current state of crop protection research. The questionnaire comprised 39 questions that were designed as multiple choice, single answer, Likert scale answers, matrix, and open-ended questions. These questions were structured to i) understand a respondent’s profile (institution represented, country, gender, age group, education level); ii) identify priority crops and biotic stresses where the respondents were working on and their perception of institutional level of expertise; iii) determine facilities and infrastructure available for crop protection research; iv) understand current extent of use of digital tools for phenotyping and limitations to adoption; v) establish capacity development needs (e.g., priority training areas) to conduct crop protection research; vi) highlight challenges/limitations faced by the researchers at their respective institution to undertake crop protection research activities. Countries were selected across Africa based on the dynamics of crop protection activities in each country. The plant health team leads or their delegates from NARES institutions filled out the survey.

### Analysis

The participating countries were further classified into East (EA), Southern (SA), Western (WA), or Central Africa (CA) based on their geographical location on the African continent. All collected data were coded and analyzed using Excel and STATA software, version 14. Descriptive statistics and comparison were used to present the data. Frequency distribution was used to analyse categorical variables such as regions, countries, gender, crop varieties, and respondents’ profiles. Cross-tabulation analysis was performed to determine the relationship between categorical variables. Qualitative variables were converted into quantitative variables to allow statistical ranking of categorical data. The Likert scale was used to quantify the perceived expertise of crop protection units on the selected crops, with scale: 1 = Excellent, 2 = Very good, 3 = Good, 4 = Fair, ranking of technical areas where capacity is needed (1 = Strongly Needed; 2 = Needed, 3 = Slightly needed, 4 = Not needed). For each country, the average expertise rating per crop and the average rating of capacity needs in technical areas were computed based on responses from individual participants. The usage of digital tools for field evaluation data and documentation was assessed by asking respondents about their familiarity with digital tools and the approaches used in data documentation. Drawbacks to the adoption of digital tools were assessed through a provided list, and respondents were able to select the most relevant responses for their institutions. The mean ranking of capacity needed on different aspects by respondents from the same institution was computed.

## Results

### Respondents’ demographics

Fifty-one (51) respondents from 36 plant health units spread across 26 countries in SSA participated in the survey. Based on the regional classification of the countries that participated in the survey, West Africa had ten; East Africa had six, while Southern Africa and Central Africa were each represented by five countries (Table 1).

**Table 1:** Distribution of respondents by countries and number of institutions across the different regions of Africa.

| African Regions | Number of participating countries | Number of institutions | Percentage of institutions | Number of respondents |
| --- | --- | --- | --- | --- |
| East Africa | 6 | 12 | 33.3 | 15 |
| Southern Africa | 5 | 7 | 19.4 | 7 |
| **Total ESA** | **11** | **19** | **52.8** | **22** |
| Central Africa | 5 | 5 | 13.9 | 12 |
| West Africa | 10 | 12 | 33.3 | 17 |
| **Total WCA** | **15** | **17** | **47.2** | **29** |
| **Grand Total** | **26** | **36** | **100** | **51** |

Consequently, West Africa was represented by the largest proportion of respondents (33%), while Southern Africa had the least (14%) (Figure 1).

**Figure 1:**
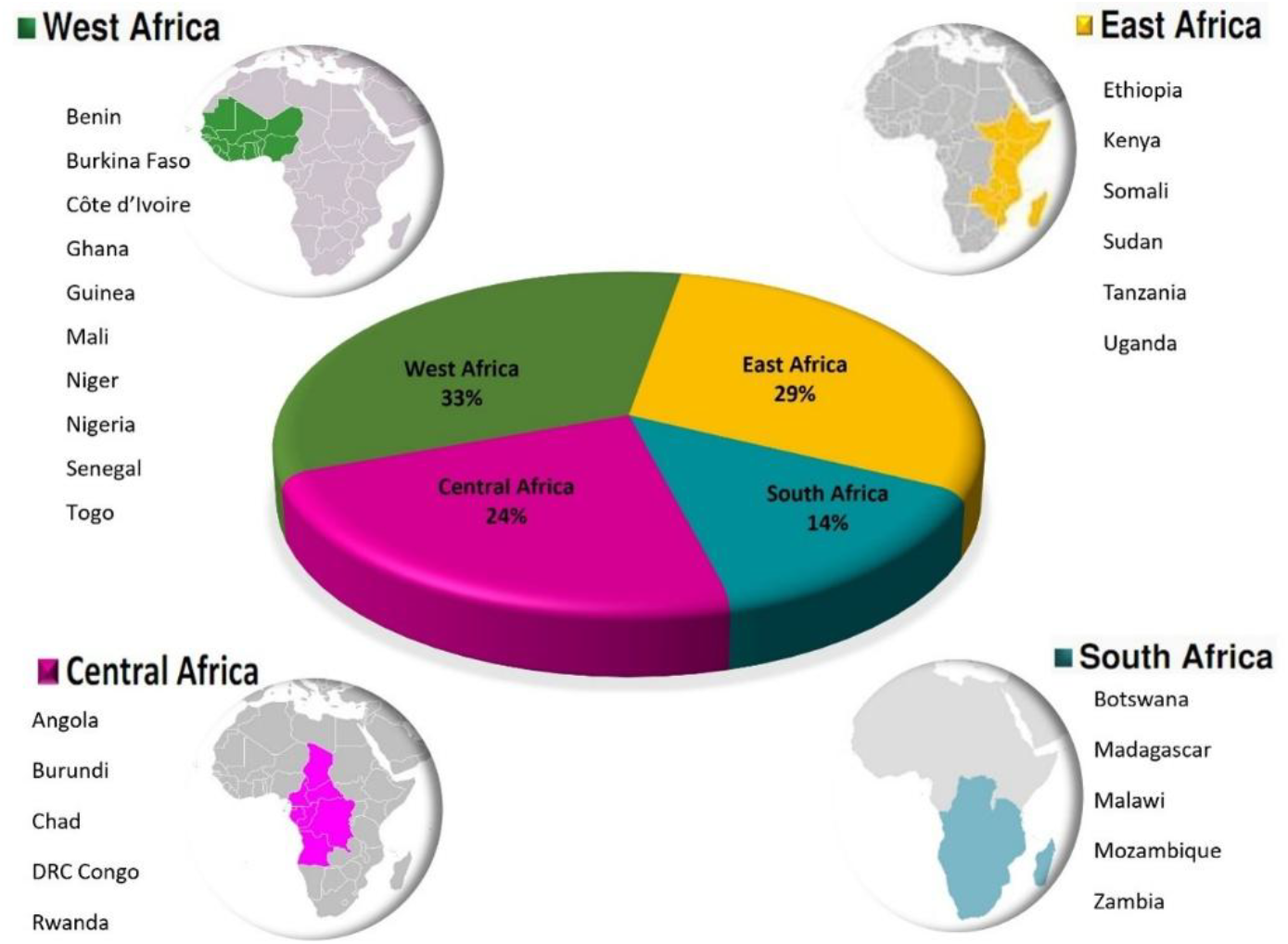
Regional distribution of survey respondents across Africa. Respondents were drawn from four sub-regions in East, Southern, Central and West Africa

Combined, 56.9% of all respondents came from West and Central Africa (WCA) regions, while East and Southern (ESA) regions had 43.1%. Out of the 36 plant health units represented in the survey, East and West Africa had majority 33.3% in and the least, 13.9%, from Central Africa (Table 1).

### A highly skilled workforce characterized by gender disparities and under-representation of younger trained staff

Forty-two (82.3%) of the respondents were male. West Africa had slightly more female respondents (29.4%) compared to the other regions (Figure 2; Supplementary 3). There was, however, no significant difference (p < 0.05; χ2 = 4.30204) in gender proportions between the four regions. The largest proportion of respondents (49%) were aged between 35 and 45 years, out of which only 2 (<1%) were female. Only five of the respondents were below 35 years old, of whom three were females and two males. All female respondents from EA and SA were below 45 years old, while none of the respondents in WA and CA were below 35 years old. Furthermore, most respondents from WCA were above 55 years (Figure 2; Supplementary 3). The distribution of respondent age groups did not, however, differ significantly (p>0.05; χ2 = 10.10) between the regions.

**Figure 2:**
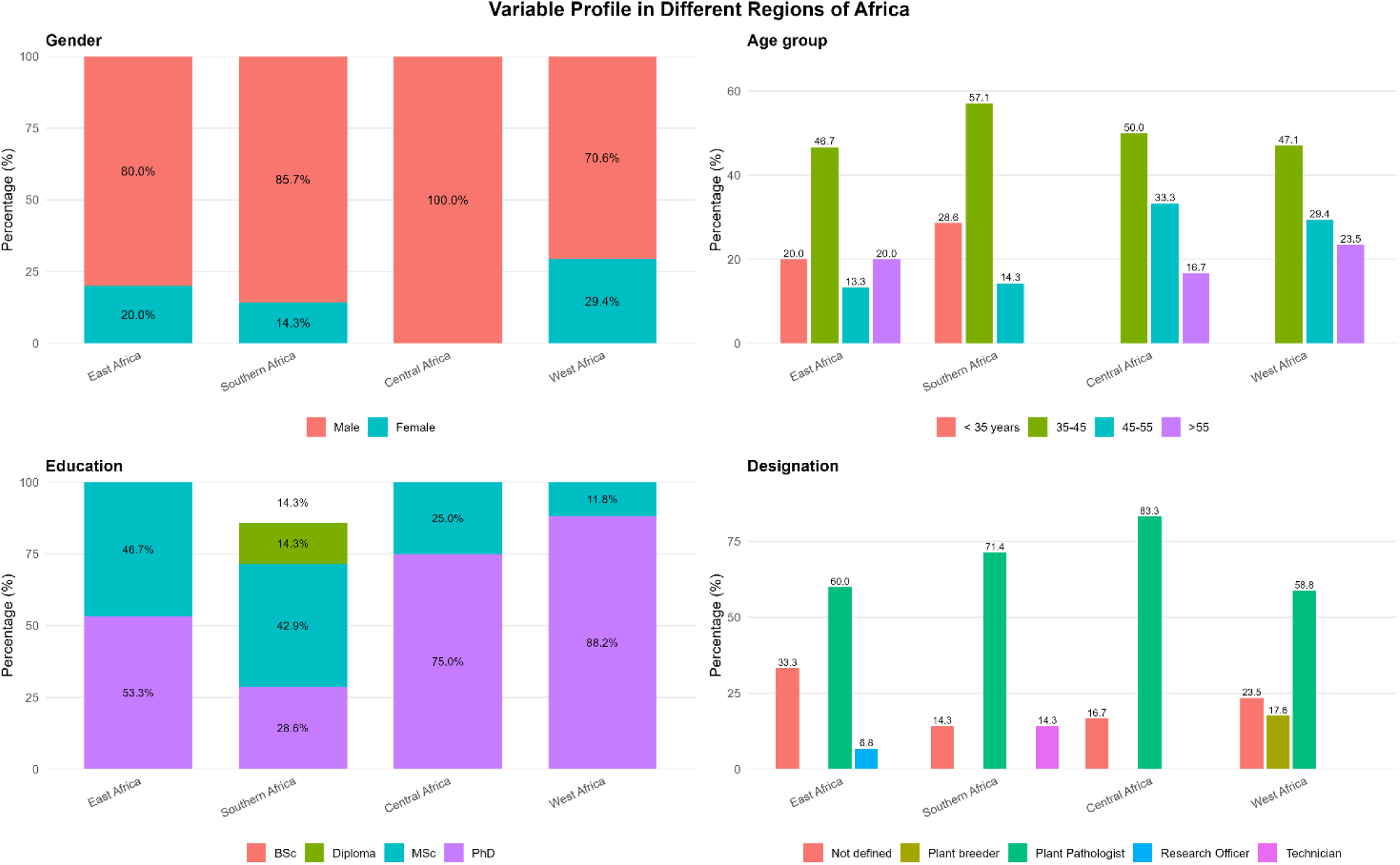
Regional distribution of survey respondents by gender, age group, education level, and professional designation across East, Southern, Central, and West Africa. Percentages indicate the proportion of respondents within each region

The largest proportion of respondents are PhD holders (N=34; 67%), followed by MSc holders (N=15; 29%). There was significant difference (p < 0.05; χ2 = 19.53) in level of education between the 4 regions. In terms of gender, only 6 (17.6%) out of the 34 PhD holders are female (Figure 2; Supplementary Table 2).

Sixty-eight percent of respondents who participated in this survey are plant pathologists working in a plant health unit of their institutes. A few breeders (18%) from West Africa and 14% of technicians from Central Africa also responded to the survey. The rest of the respondents identified themselves as technicians or research officers, while 24% from all sub-regions did not specify their area of specialization (Figure 2; Supplementary Table 2).

### Crop prioritization and distribution of plant health expertise across regions

Based on the number of respondents selecting each crop, the priority crops were ranked. Rice and maize were the most widely covered crops, with 85% and 82% of the respondents across the region working on these crops. Beans (57%), groundnut (55%), cassava (54%), banana (49%), and millet (48%) had moderate engagement with notable variation across regions. Lentil, yam (N=2; 4%), and chickpea (N=4; 8%) were represented by few respondents, showing limited geographical and institutional focus (Supplementary 4).

Respondents also ranked plant health expertise existing in their institutions for the different priority crops. The ranking of expertise varied across crops and countries. Within a country, expertise in some crops was ranked highly (Excellent), while for other crops it was good or fair. The most notable exceptions are Benin and Ghana, which ranked the expertise of all the crops as excellent. Mali, Togo and Kenya, on the other hand, ranked their expertise on all crops as either good or fair (Supplementary Table 5).

### Priority biotic stresses and screening capacity within NARES in SSA

Fungal pathogens and viruses are the priority biotic stresses in Africa, as an average of 100% and 90% of respondents, respectively, indicated that their crop protection units support their detection and screening, followed by insects at 80% and bacteria at 78%. Phytoplasma was the least studied (<10%) and is currently only detected in East and Southern African countries (Figure 3).

**Figure 3:**
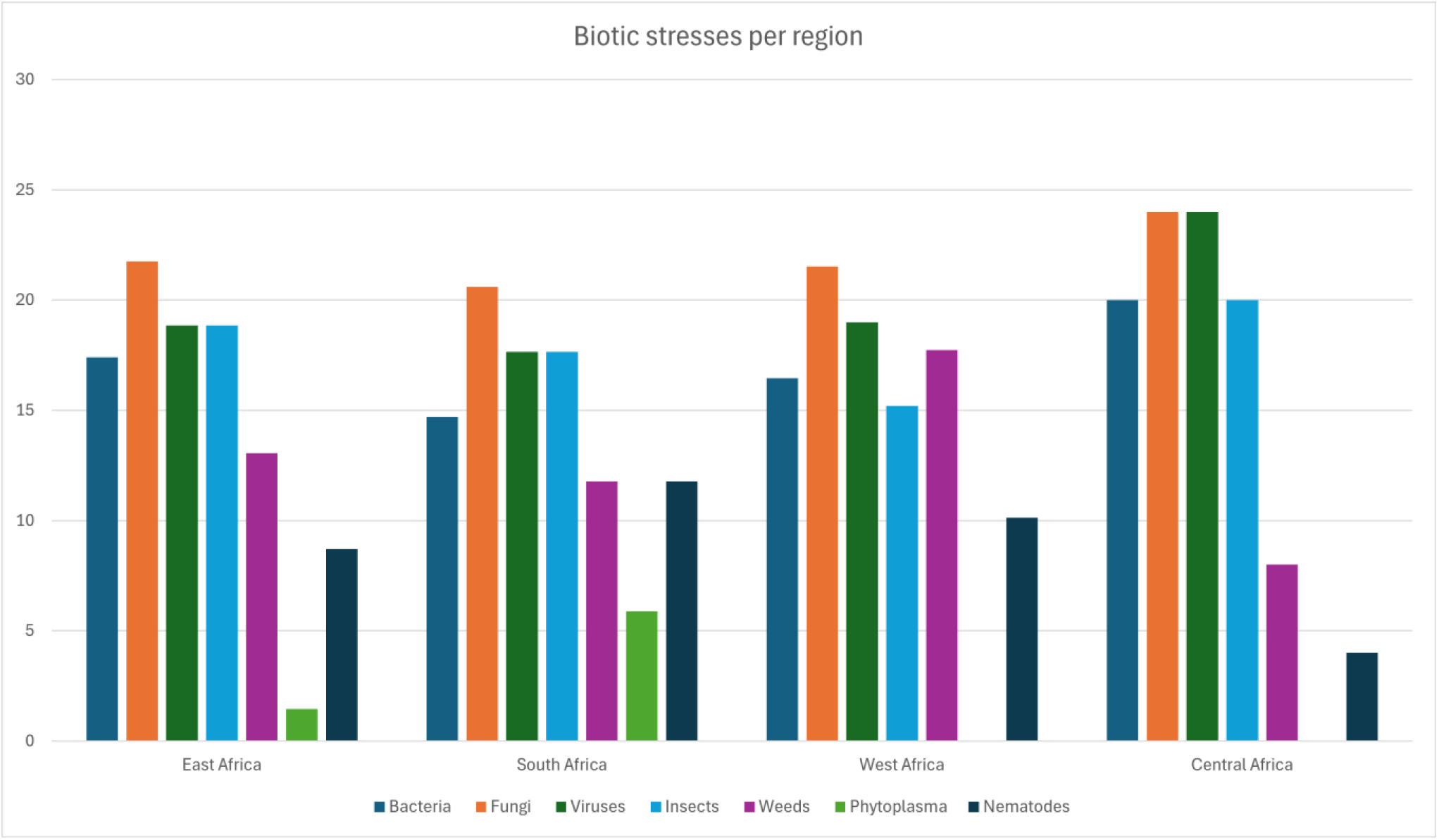
Percentage of respondents from plant health units in a region detecting different biotic stresses in each region

Although respondents indicated that their plant health units are capable of screening several specific biotic stresses, when asked to list specific stresses where technical capacity improvement was required, a number of insect pests, weeds and diseases were mentioned, including Striga weed, wild rice, fall armyworm, stem borers, *Tuta absoluta*, Fusarium wilt in peas, rice blast, powdery mildew, anthracnose, leaf spots (early and late) of groundnut, finger millet blast, banana bunchy top, Fusarium wilt TR4, and Maize lethal necrosis disease, among others (Supplementary 6). These varied across countries and appeared to be related to the crops that plant health units work on.

To further interrogate the capacity of different plant health units to phenotype against different biotic stresses, we asked the respondents to list the biotic screening infrastructure they had. Crop protection and molecular laboratories were more abundantly listed by 59% and 43% of the respondents, respectively, across the regions, while dirty plots (31%) and endemic plots (29%) were the least. The availability of screening facilities varies between regions. For example, West Africa (12) and East Africa (10) had a higher number of respondents with crop protection laboratories; East Africa has more glass/screen houses (10) while West Africa reported more dirty plots (10). Distribution of molecular labs was more uniform compared to other facilities. Southern Africa and Central Africa have fewer screening facilities (Figure 4).

**Figure 4:**
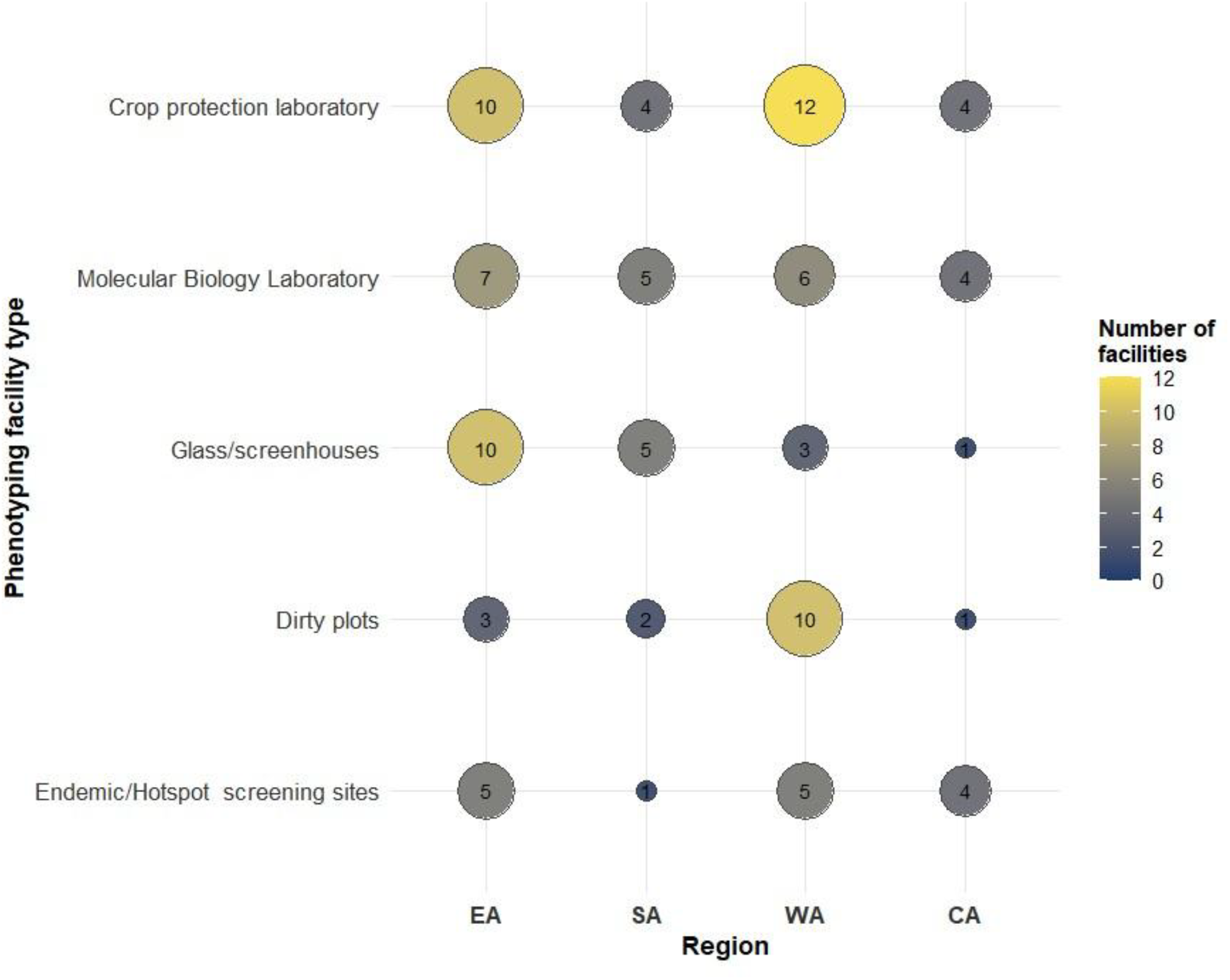
Regional distribution of phenotyping facilities for biotic stress screening in Africa. Bubble size and color indicate the number of respondents reporting each facility type across East Africa (EA), Southern Africa (SA), West Africa (WA), and Central Africa (CA); values within bubbles show the number of respondents.

### Operational fragmentation of screening capacity, lack of standardization and limited use of digital tools in the region

The result indicated access to lab equipment as the major constraint (N = 43; 84% in SSA, followed lack of trained personnel (N = 41; 80%), lack of greenhouse screening facilities (N= 40; 78%), access to protocols (N = 30; 59%), and lack of purified inoculum or germplasm checks (N= 24; 47%). Lack of endemic screening sites where biotic stresses are established, and national phytosanitary restrictions hampered only 14% and 10% of the respondents (Figure 5).

**Figure 5:**
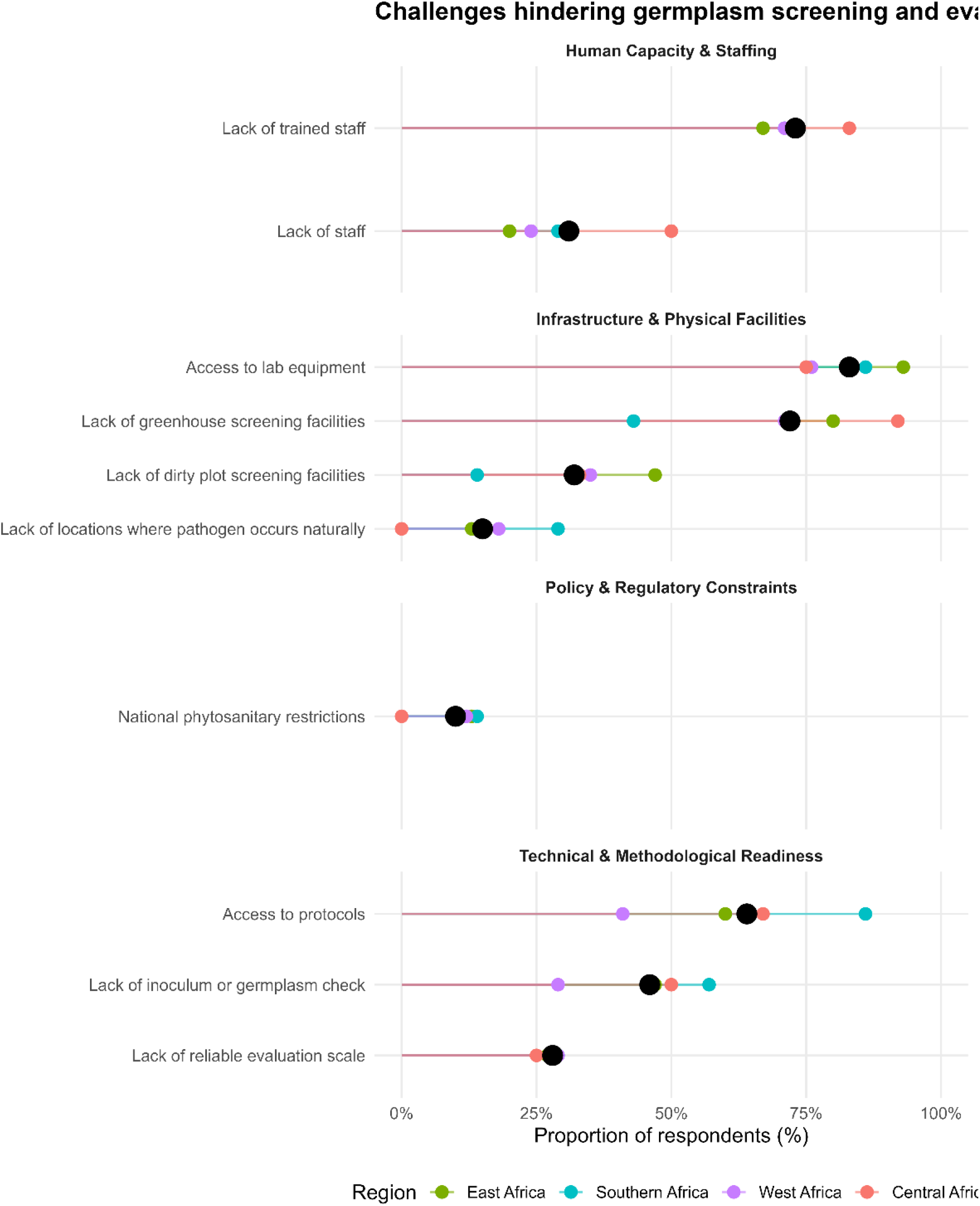
Regional variation in challenges constraining germplasm screening and evaluation for biotic stress resistance in Africa. Constraints were categorized into human capacity and staffing, infrastructure and physical facilities, policy and regulatory constraints, and technical and methodological readiness. Points represent the percentage of respondents from East, Southern, West, and Central Africa reporting each challenge, with black points showing the overall regional average. Higher percentages indicate more widely reported constraints.

Most respondents (53%) reported that their crop protection units lack standard operational procedures (SOPs) that guide evaluations for biotic stresses. In East Africa, 67% of the units have SOPs, while in Central and Southern Africa the majority (75% and 57%), respectively, lack SOPs (Figure 6).

**Figure 6.**
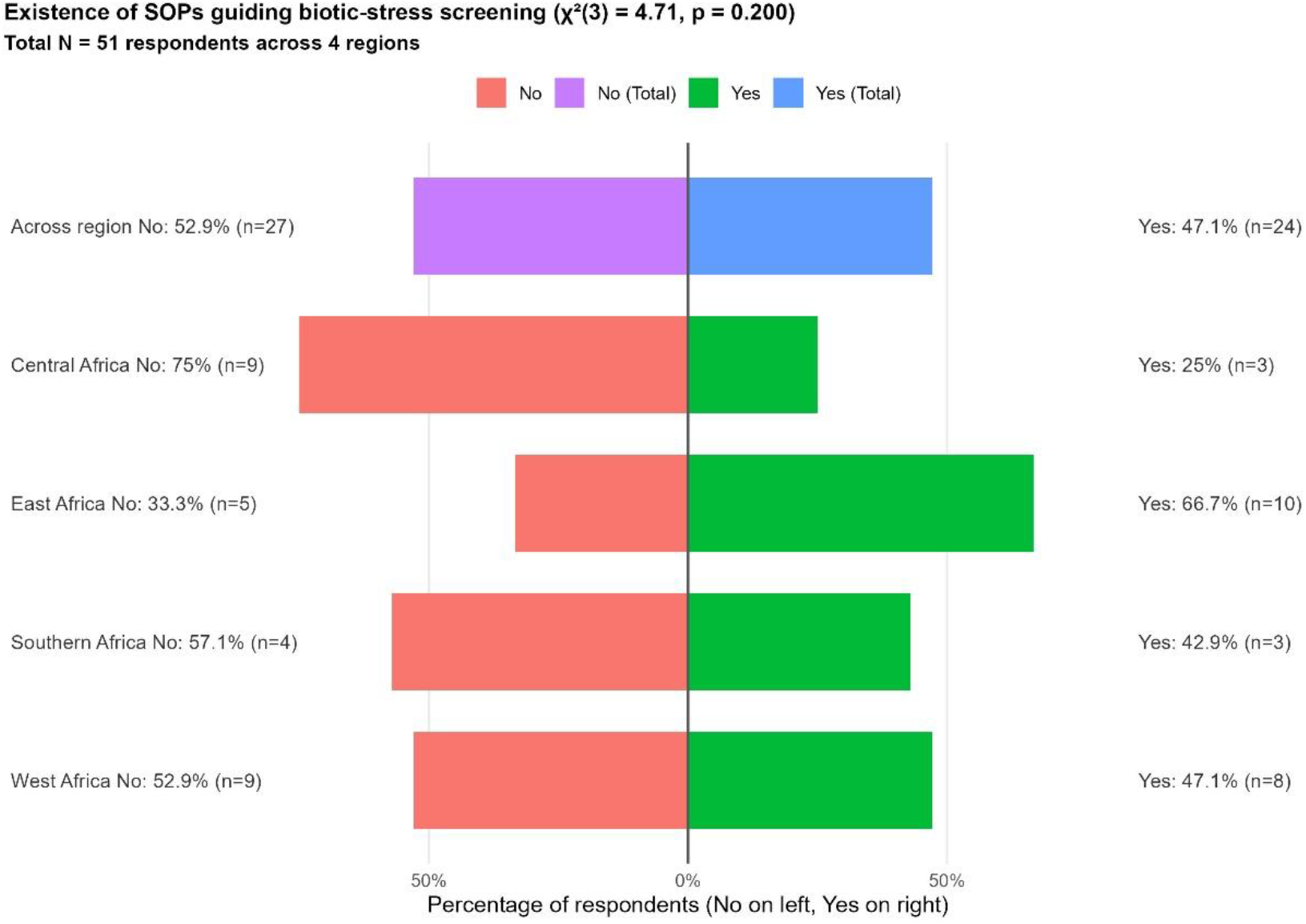
Proportion of Plant health units with established standard operating procedures guiding screening of biotic stresses in different regions

Participants in this survey were asked if they were familiar with the use of digital tools (phones or tablets) in phenotyping data documentation. Only 10% of the respondents reported using them regularly. The majority, 57%, have heard of these tools but have not used them, while 16% tried them but do not regularly use them. Twelve percent of respondents were not familiar with the use of any digital tools in biotic stress phenotyping and data documentation (Table 2).

**Table 2:** Data documentation methods and limitations to digital tools adoption by agricultural research institutions in sub-Sahara Africa.

| Phenotyping data documentation methods | Percentage of data documentation approaches in Africa |  |  |  |  |
| --- | --- | --- | --- | --- | --- |
|  | East Africa | Southern Africa | West Africa | Central Africa | Average |
| Institutional report system | 7 | 14 | 10 | 15 | 11 |
| Excel sheets within institutional computers | 7 | 29 | 10 | 19 | 14 |
| Excel sheets within personal computers | 53 | 43 | 48 | 37 | 44 |
| Notebooks | 33 | 14 | 19 | 11 | 19 |
| Institutional database | 0 | 0 | 14 | 19 | 11 |
| Limitations to adoption of digital tools | Proportion (%) of respondents reporting each limitation |  |  |  |  |
| Lack of finances | 39 | 44 | 44 | 46 | 43 |
| Limited access to internet | 18 | 25 | 30 | 27 | 25 |
| Limited skills and knowledge | 42 | 31 | 25 | 27 | 32 |

We also found out that the majority of represented institutions (77%) lack centralized documentation systems for phenotyping results like databases and institutional computers. Most of the screening results are documented in Excel files within personal computers (44%), or notebooks (19%).

In East and Southern Africa, none of the institutions reported having an institutional database for data documentation. The majority (53%) document their screening results in Excel sheets within personal PCs, and 33% in notebooks, while only 13% are institutionalized (institutional report system or Excel file within institutional PC) (Table 2). Slightly more institutions in West and Central Africa have institutionalized documentation systems. However, there is still a proportion of institutions that record their data on personal computers and notebooks (Table 2). We further wanted to understand why the use of digital tools was limited in the region. According to 43% of the respondents, lack of financial support for digitization, lack of skills and knowledge (32%) and poor internet coverage (25%) are the greatest impediments to adoption (Table 6). The limitations are similar across the different sub-regions, although internet coverage in East Africa was mentioned by a slightly lower proportion of respondents (Table 2).

To better support their institution in phenotyping biotic stresses, we asked respondents to rank different technical, research and data management aspects where capacity building is required. This information was asked for in 4 thematic areas (Figure 7A, B, C & D). Analysis shows that across thematic areas, the most urgent training needs are concentrated in research methods, data management, host-pathogen interaction, and selection of resistance improvement, all showing consistently high proportions of “highly needed” responses across all components. Data-related skills (data analysis, capture, and interpretation) and resistance breeding approaches form the highest priority cluster, indicating a strong demand for strengthening data-driven breeding and decision-making systems. The second priority tier is field diagnostics and pathogen handling, where most topics show substantial but slightly more distributed responses between “highly needed” and “needed,” suggesting these are critical operational skills that require reinforcement but are relatively better established. In contrast, pest and stress management emerge as the lowest-priority thematic area, with responses more evenly spread across “needed” and “slightly needed,” indicating that while important, these skills can be addressed in a phased manner. Overall, training investments should prioritize data-centric capabilities and resistance biology, followed by strengthening disease diagnostics systems, while pest and stress management can be scheduled as a secondary priority.

## Discussion

Until recently, comprehensive assessments of the capacity of agricultural development specialists and research facilities in sub-Saharan Africa (SSA) were limited. This constrained the ability to identify and address critical gaps in both human and institutional capacity, including deficiencies in agricultural research infrastructure (IFPRI, 2016). This study provides systematic, multi-country assessments of plant health research capacity within National Agricultural Research and Extension Systems (NARES) across sub-Saharan Africa (SSA), addressing a critical and long-standing knowledge gap. Drawing on survey data from 51 respondents across 36 plant health units in 26 countries, the findings reveal a region with great potential but facing limitations in various interconnected categories: workforce capacity, limited infrastructure, limited digital data tools adoption, gender and generation inequality and the case for a regional, coordinated investment architecture.

### A qualified but functionally constrained workforce

The study reveals the presence of a relatively well-qualified workforce with a substantial proportion of scientists holding postgraduate qualifications, indicating a strong foundation of technical expertise. However, this potential is constrained with the lack of practical experience to handle activities like germplasm evaluation and biotic stress screening. Similar constraints have been documented elsewhere, where both shortages in numbers and limited practical training opportunities restrict implementation of research activities (Boulanouar, 2013; Miller et al., 2010a).

In SSA, these limitations are systemic and widespread and get worse due to inadequate infrastructure, limited operational funding, and continued outmigration of skilled scientists from Africa (Jayne et al., 2023). Restricted and a lack of regular programs to allow access to continuous professional development further exacerbate this gap. Globally, SSA has the lowest researcher density of approximately 70 researchers per million people compared to 2,640 in North America and 4,380 in Japan (IFPRI, 2016). While increasing researcher numbers, it is important to ensure that expertise with needed specializations and experience are onboarded, enhance their retention in the system, and finally ensure the system optimally facilitates their functions.

### Inadequate infrastructure as an impediment to biotic stress screening in the region

Infrastructure deficits represent a central constraint to plant health research in SSA. Although many institutions report the ability to detect major pests and diseases, fewer than half possess essential facilities such as screenhouses, plant pathology laboratories, and managed stress environments. These findings align with earlier studies highlight persistent infrastructure gaps across SSA (Miller et al., 2009, 2010b). Such deficiencies directly limit the reliability and scalability of phenotyping, which underpins resilient resistance breeding. Without controlled and uniform screening environments, data quality is compromised, slowing genetic gain and reducing the impact of plant health in crop improvement programs. These challenges are rooted in chronic underinvestment in agricultural research and development, leaving many NARES under-resourced and unable to deliver sustained plant health research outputs (Jayne et al., 2023; Stads et al., 2022).

### Fragmented screening facilities and a lack of standardized operating procedures

Fragmentation in screening efforts further constrains plant health research; for instance, despite the low funding, there is poor coordination across institutions or countries. This forces each of the plant health facilities to operate in isolation, which limits efficiency and reduces opportunities for shared learning and resource optimization. In parallel, the absence of standardized protocols and operating procedures undermines data quality, comparability, and reproducibility. Without harmonized screening methodologies and biotic stress rating systems, collaboration is constrained, and confidence in isolated research outputs is totally reduced and incomparable, hence limiting scaling of crop improvement goods at the regional level. Standardization is therefore essential for enabling coordinated research and the development of broadly adapted resistant varieties.

### Limited modernization of data capture and management

Digitalization remains a critical but underutilized component of plant health research systems. Digital tools can enhance data accuracy, enable real-time quality control, and improve data sharing and integration (Bacco et al., 2019; Deichmann et al., 2016; Knust, 2022). However, adoption of modern tools across SSA remains low, with many researchers relying on paper-based records or unstructured personal storage systems. This limits data integrity, accessibility, and long-term usability, particularly in the context of high staff turnover. Weak data management systems constrain multi-location analysis and extrapolation. For modern data management tools to be effectively adopted, key barriers such as limited infrastructure, inadequate technical capacity, financial constraints, and unreliable internet connectivity must be addressed (Carvajal-Yepes et al., 2022; Mhlanga & Ndhlovu, 2023). Addressing these interconnected constraints is essential to enabling data-driven research and the modernization of crop improvement programs.

### Gender disparities and weak generational succession

The workforce composition revealed significant gender and generational imbalances. Women remain underrepresented, particularly in senior scientific roles, reflecting persistent structural barriers. Globally, women account for approximately 20% of agricultural researchers, and SSA reflects similar disparities (Carvajal-Yepes et al., 2022; Research Institute (IFPRI), 2016). At the same time, the limited presence of early-career scientists indicates weak succession planning within NARES. This creates a risk of knowledge loss as senior scientists retire, undermining long-term system sustainability. Addressing these imbalances requires targeted recruitment, mentorship, and institutional reforms to support equitable career progression and strengthen the pipeline of young scientists.

### Towards integrated and sustainable plant health regional capacity development in SSA

The constraints identified spanning human resources, infrastructure, digital systems, and institutional coordination are interconnected and systemic. Addressing them requires a shift from fragmented, project-based interventions towards long-term, integrated investment strategies. While international partners such as the World Bank, African Development Bank, and CGIAR have contributed to capacity building, these efforts have often been time-bound and commodity-specific, limiting sustained institutional impact (Jayne et al., 2023; IFPRI, 2016). Regional collaboration supported by governments offers a viable pathway to overcome resource limitations and fragmentation. Shared infrastructure, harmonized protocols, and coordinated training can improve efficiency and reduce duplication (Jayne et al., 2023; Stads et al., 2022). Emerging initiatives such as the African Dryland Crop Improvement Network (ADCIN) demonstrate the potential of such coordinated, multidisciplinary approaches. Given the complexity of agricultural systems in SSA characterized by diverse agroecologies and multiple interacting constraints, including pests, diseases, soil fertility decline, and climate variability (Carvajal-Yepes et al., 2022; Neuenschwander et al., 2023; Savary et al., 2017), a systems approach to capacity development is essential. Strengthening plant health research will require aligned investments in infrastructure, workforce development, digitalization, and regional integration to ensure that existing scientific capacity can be effectively translated into improved agricultural productivity and food security.

### Limitations and future directions

This study is subject to some limitations. Findings are based on self-reported data, which may introduce bias. Regional representation is uneven, with West Africa overrepresented, potentially affecting comparative analyses. The cross-sectional design limits inference on temporal trends and causality, and the absence of financial data constrains direct linkage between investment levels and capacity outcomes. Future research should prioritize longitudinal analyses to assess the impact of capacity investments over time, as well as cost-effectiveness comparisons across intervention models. Complementary qualitative studies on institutional governance, culture, and retention dynamics would deepen understanding of systemic constraints. Expanding geographic coverage will further enhance the robustness and generalizability of findings.

## Compliance with ethical standards

All participants in the survey voluntarily consented to participate in this survey.

## Acknowledgements

Authors are grateful to all respondents who took part in the online survey. The views expressed in this manuscript are those of the authors and do not represent views of affiliated organizations and the funding agencies.

## Author Contributions

HG conceptualized the study and guided implementation, and MS writing. AEA designed the study and participated in writing and editing the MS, BS conducted the online survey, collected data and drafted survey report, JNK data curation, interpretation, led in drafting of first and final MS, AR, RD, HS supported in data visualization and MS review, NB, DI, JB, AP and TMN reviewed and edited the various versions of the MS. All authors read and approved the final manuscript.

