## Supplementary 6 for "Advancing Plant Health in Sub-Saharan Africa: Understanding National Agricultural Research Institutions capacities for Strategic Investment Priorities to Boost Genetic Gains Against Crop Biotic Stress"

Supplementary 6: Priority biotic stresses for capacity building in the region

| **Region** | **Country** | **Weeds** | **Insect** | **Pathogens** |
| --- | --- | --- | --- | --- |
| **East Africa** | Ethiopia |  |  | Pulse viruses, Pulse root diseases, wheat rust |
|  | Kenya | couch grass, sedges; weeds: parasitic weeds: striga in cereal crops & alectra weed in cowpea | Pod borer of pigeon pea and fall armyworm of sorghum; pests of legumes, aphids on legumes, thrips in legumes, sorghum shoot; fly, fall army worm in sorghum, stem borer in sorghum; storage pests of pigeon peas, mung beans, sorghum, millets, cassava, groundnuts, and chickpea | Potato wilt and potato cyst nematodes, fusarium wilt and Ascochyta blight of chickpea, rice blast, finger millet blight; c. Bacterial diseases: common bacterial blight of beans and mung beans in legumes, pod sucking Fungal diseases: Fusarium wilt of pigeon pea, Ergot in sorghum and millet, anthracnose of beans, mung beans, sorghum, powdery mildew, Viral diseases: viral diseases of beans, mung beans, cassava and sweet potato |
|  | Tanzania | Striga weeds  *Oryza longistaminata*, wild rice | white flies, aphids, Fall armyworm | Fusarium wilt, Sweetpotato virus disease (SPVD), cassava mosaic disease (CMD); *Fusarium oxysporum* in cotton crop, Cassava Broun Streak Virus, Cassava Mosaic Virus, *Xanthomonas oryzae* pv. oryzae, Rice Yellow Mottle Virus; Cassava brown streak Cassava mosaic virus Powdery mildew disease Late and early leaf spot Blight disease on cashew, blast, RYMV, BLB, *Pyricularia oryzae* |
|  | Uganda | Striga | Midge; maize fall armyworm pest | Anthracnose, Bacterial blight, Cassava Brown Streak Disease virus, Groundnut rosette virus, Citrus leaf and fruit spot disease pathogen, Pigeonpea wilt pathogen  Finger millet Blast, Citrus leaf and fruit spot disease, pigeonpea wilt pathogen & groundnut rosette |
|  | Sudan | *Striga hermonthica ; Cyperus rotundus; Cynodon dactylon* | Stem borer; Fall armyworm; Sorghum midge; African bollworm; Head bug, Shoot fly | *Sphacelotheca reiliana*; cercosporin leaf spot; helminthospora |
|  | Somalia | *Sonchus exsauriculatus ; Sorghum arundinaceum ; Tetrapogan tenellus; Triantherma pentandra*; Solanum spp | *Chilo partellus*, *Cylas puncticollis* | *Rhopalosiphum maidis*; *Antigastra catalaunalis* ; *Sporisorium sorghi* |
| **Southern Africa** | Botswana | Striga spp, Electra spp, | Fall armyworm, Diamond Back Moth, Fruit fly (bactocera), quelea birds |  |
|  | Madagascar |  | Grubs, Nematodes, *Tuta absoluta* | Erwinia sp., *Aspergillus flavus*, vanilla Fusarium wilt, Foc TR4, CBSD, Dickeya, *Magnaporthe grisea*, bean fat, |
|  | Malawi | *Striga asiatica* (witchweed), *Elusine indica* | Fall armyworm; *Tuta absoluta* | Banana bunchy top Rice yellow mottle, Cassava mossac virus; *Fusarium oxysporium* TR4 (Fusarium wilt of banana), Maize Lethal Necrosis disease, Rice blast |
|  | Mozambique | Red rice, cyperus ; rotundus, cinodon ; dactolon, Red rice | Thacth borer | Fusarium wilt FOC TR4, Cassava brown streak vírus blast and mosaic, Maize Lethal yellow disease and Rosette, Bean Golden mosaic and Coconut lethal yellow disease, blast |
|  | Zambia | *Salvinia molesta*, *Mimosa pigra*, Cyprium spp, itch grass and *Cynodon dactylon* | Pests: Fall armyworm, *Tuta* *absoluta*, fruit flies | Rice yellow mottle virus, cassava brown streak virus, Maize lethal necrosis disease, Avocado sunblotch virus, maize chlorotic mottle virus |
| **West Africa** | Benin | *Striga hermonthica*  *Imperata cylindrica*  *Cynodon dactylon*  *Cyperus haspan*  *Killina bulbosa* | The armyworm  Cashew stem borers  Knot nematodes  Tomato cutworm  The whitefly and Fruit fly | The crown rot of vegetable and fruit crops  Citrus fungal diseases  Fungal diseases of the cashew tree  Banana bunchy top disease |
|  | Burkina Faso | *Commelina benghalensis* L.; *Ipomoea eriocarpa* , *Ottboellia cochinchinensis*; *Striga hermontica*, Cyperus sp. | *Chilo zacconius*, *Orseolia oryzivora*, *Bemisia tabaci*, *Helicoverpa armigera*, grain storage pests | Fungi (*Fusarium* spp, Botryodiplodia spp, Phoma spp; Botrytis spp, *Magnaporthe grisea*), Bacteria (Xanthomonas spp, *Ralstonia solanacearum*, Pectobacterium spp; Erwinia spp, Pseudomonas spp), Imperata yellow mottle virus (IYMV), Maize yellow mosaic virus (MaYMV), Banana streak virus (BSV), Rice necrosis virus (RSNV), sorghum mosaic virus (SrMV) sugarcane mosaic virus (SCMV), Peanut clump virus (PCV), (Maize yellow dwarf virus-RMV2 |
|  | Cote d'Ivoir |  |  | Pantoea, *Sarocladium oryzae*, *Fusarium fujikuroi* *Bipolaris oryzae* |
|  | Ghana | parasitic weeds (*Striga*) | Sorghum midge, cowpea thrips, Parasitic nematodes, stem borer of sorghum and pearl millet, head miners of sorghum and millet | Rice blast Grain mold of sorghum, Downy mildew of pearl millet, late and early leaf spot of groundnut, Groundnut rust, sorghum and millet smut, sorghum anthracnose, macrophomina rot, ergot of sorghum and pearl millet |
|  | Guinea | *Cyperus rotundus, Biden pilosa* | Corn armyworm | *Ustilago maydis*, verticilian wilt. |
|  | Mali | Wild rice (*Oryza longistaminata* and *Oryza bartihii*),  *Echinocloa colona .Iscamum rigosum* | rats, seed-eating birds | Viruses (RYMV), bacteriosis (BLB) |
|  | Níger | Striga spp, *Echinochola* spp, Broomrape, Dodder, Cyperus spp, | *Spodoptera frugiperda*, Norda blitelis, *Contarina sorghicola*, *Heliochelus albipunctela*, *Tuta absoluta*, Root-knot nematodes | Bacterial leaf blight, *Ralstonia solanacearum*; *Xanthomonas campestris*; Rice Yellow Motttle, *Pectobacterium carotovorum*, |
|  | Nigeria | Striga, | Armyworm, stem borer | *Aspergillus, Colletotrichum, Sclerospora , Phythopthora, Fusarium, Altenaria* |
|  | Senegal | Striga, sedge, grasses, | caterpillars, stem borers, sucking borers, defoliators | BLB, RYMV, pyricularios, downy mildew, anthracnose |
|  | Togo | Striga, | pests (armyworm, stem and ear borers, pod borers, stock | Legume viruses (rosette, mosaics), cereal viruses (MSV, MaYMV, SCMV), leaf spot diseases (anthrachnose, downy mildew, leaf spots, Sigatoka) |
| **Central Africa** | Angola |  | Fall armyworm, fruit fly, Green Mite and Mealybugs, Birds | Rice blast, Maize Lethal Necrosis Disease, Fusarium wilt of banana, Banana bunch top virus, Cercospora *angolensis*, Late blight and Early blight of potatoes  Beans: Bacterial blight; Bacterial brown spot, Bean Common Mosaic Virus  Cassava: Cassava Brown Leaf Spot, cassava brown streak virus, Bacterial Blight, Brown spot |
|  | Burundi |  |  | Rice Yellow Mottle Virus, Bean: Damping-off and leaf angular spot, Cassava: Cassava brown streak virus, Maize: Maize Lethal Necrosis viruses |
|  | Chad | weeds, | Desert locust, red locust, grasshoppers, seed-eating birds, termites, rodents | Fungi, viruses, bacteria |
|  | DRC Congo |  | *Acraea acerata,* *Spodoptera frugiperda* Prostephanus, Antestiopsis, *Bemisia tabaci* | Fusarium wilt, Blaster, Brown stripe blotch, Anthracnose |
|  | Rwanda |  |  | Rice blast |
