## Supplementary 3 for "Advancing Plant Health in Sub-Saharan Africa: Understanding National Agricultural Research Institutions capacities for Strategic Investment Priorities to Boost Genetic Gains Against Crop Biotic Stress"

**Supplementary 3:** Demographic and professional characteristics of respondents across East, Southern, Central, and West Africa. The table summarizes respondent distribution by gender, age group, education level, and professional designation, together with overall percentages and regional comparisons using chi-square (χ²) tests. Most respondents were male (84.1%), aged 35–45 years (50.8%), and highly educated, with PhD holders comprising 61.3% of participants. Plant pathologists represented most respondents (68.4%). Significant regional differences were observed only for education level (p = 0.021), while gender, age group, and designation did not differ significantly among regions (p > 0.05).

|  |  | **Variable Profile in different regions of Africa** | | | | | | | | **Overall variable summary and analysis** | | | |
| --- | --- | --- | --- | --- | --- | --- | --- | --- | --- | --- | --- | --- | --- |
|  |  | **East Africa** | | **Southern Africa** | | **Central Africa** | | **West Africa** | |  | | | |
| **Variable** | **Category** | **Total Respondents** | **Percentage** | **Total Respondents** | **Percentage** | **Total Respondents** | **Percentage** | **Total Respondents** | **Percentage** | **Percentage Mean** | **DF** | **X^2^** | **P-values** |
| Gender | Male | 12 | 80 | 6 | 85.7 | 12 | 100 | 12 | 70.6 | 84.1 | 3 | 4.3 | 0.23064 |
|  | Female | 3 | 20 | 1 | 14.3 | 0 | 0 | 5 | 29.4 | 15.9 |  |  |  |
| Age group | < 35 years | 3 | 20 | 2 | 28.6 | 0 | 0 | 0 | 0 | 12.1 | 9 | 10.1 | 0.34240 |
|  | 35-45 | 7 | 46.7 | 4 | 57.1 | 6 | 50 | 8 | 47.1 | 50.8 |  |  |  |
|  | 45-55 | 2 | 13.3 | 1 | 14.3 | 4 | 33.3 | 5 | 29.4 | 22.6 |  |  |  |
|  | >55 | 3 | 20 | 0 | 0 | 2 | 16.7 | 4 | 23.5 | 15 |  |  |  |
| Education | BSc | 0 | 0 | 1 | 14.3 | 0 | 0 | 0 | 0 | 3.6 | 9 | 19.5 | 0.02099 |
|  | Diploma | 0 | 0 | 1 | 14.3 | 0 | 0 | 0 | 0 | 3.6 |  |  |  |
|  | MSc | 7 | 46.7 | 3 | 42.9 | 3 | 25 | 2 | 11.8 | 31.6 |  |  |  |
|  | PhD | 8 | 53.3 | 2 | 28.6 | 9 | 75 | 15 | 88.2 | 61.3 |  |  |  |
| Designation | Not defined | 5 | 33.3 | 1 | 14.3 | 2 | 16.7 | 4 | 23.5 | 22 | 12 | 16.6 | 0.16635 |
|  | Plant breeder | 0 | 0 | 0 | 0 | 0 | 0 | 3 | 17.6 | 4.4 |  |  |  |
|  | Plant Pathologist | 9 | 60 | 5 | 71.4 | 10 | 83.3 | 10 | 58.8 | 68.4 |  |  |  |
|  | Research Officer | 1 | 6.8 | 0 | 0 | 0 | 0 | 0 | 0 | 1.7 |  |  |  |
|  | Technician | 0 | 0 | 1 | 14.3 | 0 | 0 | 0 | 0 | 3.6 |  |  |  |
