## Supplementary 1 for "Advancing Plant Health in Sub-Saharan Africa: Understanding National Agricultural Research Institutions capacities for Strategic Investment Priorities to Boost Genetic Gains Against Crop Biotic Stress"

***SURVEY FOR AGRICULTURAL RESEARCH CENTERS/ INSTITUTIONS IN SUB-SAHARAN AFRICA***

**** SURVEY NOTES AND INSTRUCTIONS**

*** ATTENTION (!!!), IMPORTANT NOTES:**

Welcome to the Crop Protection survey and thank you for taking the time to participate! We appreciate your time and valuable input. Your responses are very important to us and will help us to improve plant health activities in the region. This Initiative aims to know the most important diseases/pests on different crops in Sub Saharan Africa and to capture the needs for capacity building in national research institutions.

**Note 1**: The survey consists of a series of questions that should only take a few minutes to complete.  Please take your time to read each question carefully, answer each question honestly. In order to cover the maximum number of crops, it is advised to work closely with the Pathology section of your National Agricultural Research Institution and send one set of answers covering the whole country.

**Note 2**: If you are out of time, you have the possibility to save your survey draft, and reopen it later and continue to respond. For this, please see the « **SAVE DRAFT**» **BUTTON**. To reopen the survey, you just have to click on the link sent to you and continue to answer the question where you stopped. Once all questions are completely answered, save your draft and then sent it now into the system by clicking on the button **SUBMIT** at the end of the questionary. Your answers will remain confidential and will only be used for research purposes.

Thank you again for taking the time to complete our survey.

**1. Which country are you from?**

*Mark only one oval.

- Angola
- Benin
- Botswana
- Burkina Faso
- Burundi
- Cameroon
- Central Africa Republic
- Chad
- Congo (Brazaville)
- Equatorial Guinea
- Eritrea
- Ethiopia
- Gabon
- Gambia
- Ghana
- Guinea
- Guinea-Bissau
- Ivory Coast
- Kenya
- Liberia
- Madagascar
- Mali
- Namibia
- Niger
- Nigeria
- Rwanda
- Senegal
- Sierra Leone
- Somalie
- Soudan
- South Soudan
- Tanzania
- Togo
- Uganda
- Zambia
- Zanzibar
- Zimbabwe
- Ohers

**2. Name of the institution / Research organization.--------------------------------------------------------------------------------------------------------------------------------------------------------------------------------------------------------------------------------------------------------------------------------------------------------------------------------------------------------------------------------------------------------------------------------------------------------------------------------------------------------------------**

.

**3. Physical address of the institution/ Center in the country:** * --------------------------------------------------------------------------------------------------------------------------------------------------------------------------------------------------------------------------------------------------------------------

**4.  Telephone and email contacts of the institution/ Center: ------------------------------------------------------------------------------------------------------------------------------------------------------------------------------------------------------------------------------------------------------------------------**

**5. Name, telephone and email contacts of the Head /leader of the organization:** *

----------------------------------------------------------------------------------------------------------------------------------------------------------------------------------------------------------------------------------

**6. Name, telephone and email contacts of the    CPCoP - SSA person from the institution/center:**  ---------------------------------------------------------------------------------------------------------------------------------------------------------------------------------------------------------

**7.  Name, title and contacts (if you are not the CP-Cop contact for your organization)**

----------------------------------------------------------------------------------------------------------------------------------------------------------------------------------------------------------------------------------

**8. What is your gender** *

Mark only one oval.

- Female
- Male
- Other
- Other:

**9. Age group**
* Mark only one oval.

- Below 35 years old
- 35-45 years old
- 45-55 years old
- Above 55 years old
- Others

**10. Level of education**

*Mark only one oval.

- PhD
- MSc
- BSc
- Diploma
- Certificate
- Other

**11. Does your Institution/Center have a Crop Protection unit or team?**

*Mark only one oval.

- Yes
- No
- hybrid unit (if integrated with another program)

**12. If YES, answer following questions:**

**Select different crops on which Protection unit or team is working.**

*Check all that apply.

- Banana
- Beans
- Cassava
- Chicpea
- Cowpea
- Groundnut
- Lentil
- Maize
- Millet
- Pigeon pea
- Potatoes
- Rice
- Sesame
- Sorghum
- Sweet Potatoes
- wheat
- Yams
- Other

**13. Level of crop protection expertise for different crops selected**

Mark only one oval per row.

|  | Excellent | Very good | Good | Fair |
| --- | --- | --- | --- | --- |
| Banana |  |  |  |  |
| Beans |  |  |  |  |
| Cassava |  |  |  |  |
| Chicpea |  |  |  |  |
| Cowpea |  |  |  |  |
| Groundnut |  |  |  |  |
| Lentil |  |  |  |  |
| Maize |  |  |  |  |
| Millet |  |  |  |  |
| Pigeon pea |  |  |  |  |
| Potatoes |  |  |  |  |
| Rice |  |  |  |  |
| Sesame |  |  |  |  |
| Sorghum |  |  |  |  |
| Sweet Potatoes |  |  |  |  |
| Yams |  |  |  |  |
| Wheat |  |  |  |  |
| Others |  |  |  |  |

**14.Please state the type of weeds, pest, pathogens that you are mostly detecting or screening:**

*Check all that apply.

- Viruses
- Bacteria
- Fungi
- Phytoplasma
- Nematode
- Insects
- Parasitic weeds
- Others

**15.  What facilities does the Crop Protection unit or team at your organization have?**

*Check all that apply.

- Crop protection laboratory
- Molecular Biology laboratory
- Screen/glass house
- Dirty plots
- Endemic screening sites
- Others

**16. Give the list of diseases/pests for which you are able to do artificial inoculations in terms of human resources and facilities------------------------------------------------------------------------------------------------------------------------------------------------------------------------------------------------------------------------------------------------------------------------------------------------**

**17. Staff working in the Crop Protection unit: How many PhD, MSc, BSc, Technicians you have in your Crop Protection unit?**

|  | 1 | 2 | 3 | 4 | 5 | 6 | 7 |
| --- | --- | --- | --- | --- | --- | --- | --- |
| PhD |  |  |  |  |  |  |  |
| MSc |  |  |  |  |  |  |  |
| BSc |  |  |  |  |  |  |  |

**18.** * **Please list top 5 priority weeds, pest/pathogens that your institution considers relevant for improving knowledge capacity for laboratory, glass house and field germplasm screening?** ---------------------------------------------------------------------------------------------------------------------------------------------------------------------------------------------------------------------------------------------------------------------------------------------------------------------

**19.** ***Is the Crop Protection Department in your country working on all prioritized pests, diseases and weeds listed in question 18? If not, please indicate others which need more work**? ------------------------------------------------------------------------------------------------------------------------------------------------------------------------------------------------------------------------------------------------------------------------------------------------------------------------------------------

**20. What are the specific hotspot regions in the country for screening the priority weeds, pest and diseases (if known)--------------------------------------------------------------------------------------------------------------------------------------------------------------------------------------------------------------------------------------------------------------------------------------------------------------**

**21. What are the Resistant, tolerant and susceptible check varieties for priority weeds, pest and diseases (if known)---------------------------------------------------------------------------------------------------------------------------------------------------------------------------------------------------------------------------------------------------------------------------------------------------------**

**22. In which area does your Crop Protection unit/ team consider to be a priority for training /capacity building**

**List of technical area**

|  | **Strongly needed** | **Needed** | **Slightly needed** | **Not needed** |
| --- | --- | --- | --- | --- |
| Identification and characterization od diseases in field |  |  |  |  |
| Collection and preparation of samples for analysis and shipment |  |  |  |  |
| Diseases evaluation in fields |  |  |  |  |
| Insect rearing and evaluation under controlled and field conditions |  |  |  |  |
| Diseases inoculation and evaluation in controlled conditions |  |  |  |  |
| Mapping and monotoring of disease incidences |  |  |  |  |
| Pest scouting and control |  |  |  |  |
| Isolation of plant pathogens |  |  |  |  |
| Plant-Fungi interactions |  |  |  |  |
| Plant-Bacteria interactions |  |  |  |  |
| Plant-Virus interactions |  |  |  |  |
| Abiotic stresses |  |  |  |  |
| Resistance genes introgression into lines |  |  |  |  |
| Seed dressing and fumigation |  |  |  |  |

**23. What are the major challenges for performing germplasm evaluation of priority weeds, pests and diseases (*if known*)**

* Check all that apply.

- Access to Protocols
- Access to lab equipments
- Lack of greenhouse screening facilities
- Lack of dirty plot screening facilities
- Lack of locations where pathogen is established occurs naturally
- National phytosanitary restrictions
- Lack of inoculum or germplasm checks
- Lack of staff
- lack of reliable evaluation scale
- Having trained staff
- Others

**24. Is there any quality system which may include standard operational procedures (SOPs) /Protocols that regulate weeds, pest and pathogen evaluation processes in your institutions?**
*Mark only one oval.

- Yes
- No

**25. How familiar are you with digital tools for documenting phenotyping data in field evaluation of priority weeds, pests and diseases?**

*Mark only one oval.

- Not at all
- I have heard of it, but never used
- I tried but it is not my major detection method
- I use it in my professional work
- Others

**26. Does your institution use any digital tools (e.g., Smartphone Apps, tablets) as a major method for field evaluation of weeds, pest and diseases?**
*Mark only one oval.

- Yes
- No
- I don't know
- I am not involved in the evaluation
- Others

**27. if YES, which one(s)**

*Mark only one oval.

- Smartphone apps
- Tablet
- Other

**28. How does your institution document germplasm screeing results?**

*Mark only one oval.

- In notebooks
- In excel sheets within personal PC
- In excel sheets within an institutional PC
- Within an institutional database
- In an institutional report system
- Other

**29. In addition to recording germplasm screening results (resistant, susceptible, tolerant) which other additional information is captured?**
*Check all that apply.

- Sowing date
- Evaluation site-name of the region
- Evaluation site-geolocation coordinates
- Aim of the trial
- Evaluation date
- Description of symptoms
- Crop season
- Vegetative stage
- Method(s) used
- Results date
- Name of evaluator
- Signature of the head crop protection
- Other

**30. What are the challenges hindering your adoption or use of digital tools?** *

*Check all that apply.

- Lack of financial support for digitalization 2
- Limited internet access
- Lack of skills and knowledge
- Not interested, not willing to use
- Other

31.

**31. Capacity rating in research methods and data management**
*Mark only one oval per row.

|  | **Strongly needed** | **Needed** | **Slightly needed** | **Not needed** |
| --- | --- | --- | --- | --- |
| Experimental design and layout |  |  |  |  |
| Data capture tools |  |  |  |  |
| Data analysis |  |  |  |  |
| Data interpretation |  |  |  |  |

**32. Are you currently part of a national or regional network for evaluating weed, pests, and disease**
*Mark only one oval.

- Yes
- No

**33.** ***If YES, what is the main purpose of this network? ----------------------------------------------------------------------------------------------------------------------------------------------------------------------------------------------------------------------------------------------------------------------------**

**34.** ***What are your expectations when your institution joins a regional and global weed, pest and pathogen evaluation network? ---------------------------------------------------------------------------------------------------------------------------------------------------------------------------------------------------------------------------------------------------------------------------------------------**

**35. Has your institution interacted with any of the CGIAR plant health scientist in your region?**
*Mark only one oval.

- Yes
- No

**36. If NO, can you please specify the reason:**

*Mark only one oval.

- We didn't know about the CGIAR
- We didn't know who to contact
- It just happens
- We are not interested
- Other

**37. If YES, Please mark which CGIAR had you contacted**

*Check all that apply.

- AfricaRice
- Bioversity
- CIAT
- CIMMYT
- CIP
- ICARDA
- ICRISAT
- IITA
- ILRI
- IRRI

**38. If YES in question 37, was this contact related to:**

*Check all that apply.

- Diagnostic services
- Disease surveys
- Sharing protocols or materials
- Requestion training
- Pathogen eradication
- Other

**39.  Any additional information you would like to be included in this surveys: ----------------------------------------------------------------------------------------------------------------------------------------------------------------------------------------------------------------------------------------------------------------------------------------------------------------------------------------------------------------**

***THANK YOU FOR YOUR PARTICIPATION***
