## Supplementary 5 for "Advancing Plant Health in Sub-Saharan Africa: Understanding National Agricultural Research Institutions capacities for Strategic Investment Priorities to Boost Genetic Gains Against Crop Biotic Stress"

**Supplementary table 5**: Respondents’ assessment of institutional plant health expertise across major crops and countries in Africa.
Scores represent perceived institutional expertise in plant health for each crop: 1 = Excellent, 2 = Very good, 3 = Good, 4 = Fair; “–” indicates crop not reported or not applicable. This table summarizes respondents’ perceptions of institutional capacity in plant health expertise across key crops in East, Southern, Central, and West Africa. Scores reflect variation in perceived expertise both within and between regions and crops, with generally stronger ratings (lower scores) observed for staple crops such as maize and rice in several countries, and more variable or weaker ratings for some legumes and root/tuber crops. The data highlight clear heterogeneity in institutional capacity across crops and geographies, suggesting uneven specialization and potential gaps in plant health support for less prioritized crops in several national programs.

| **Region** | **Country** | **Banana** | **Beans** | **Cassava** | **Chickpea** | **Cowpea** | **Groundnut** | **Lentil** | **Maize** | **Millet** | **Pigeon pea** | **Potatoes** | **Rice** | **Sesame** | **Sorghum** | **Sweet Potatoes** | **Yams** | **Wheat** |
| --- | --- | --- | --- | --- | --- | --- | --- | --- | --- | --- | --- | --- | --- | --- | --- | --- | --- | --- |
| East Africa | Ethiopia | 3 | 2 | 3 | 2 | 3 | 3 | 2 | 2 | 3 | 3 | 2 | 3 | 3 | 3 | 3 | - | 1 |
|  | Kenya | - | 3 | 3 | 4 | 3 | 4 | - | 4 | 4 | 4 | - | 4 | - | 3 | 4 | - | 2 |
|  | Somali | 2 | - | 3 | - | 3 | 3 | - | 2 | 3 | - | - | 3 | 2 | 2 | 3 | - | - |
|  | Sudan | - | - | - | - | - | - | - | 1 | 2 | - | - | - | - | 1 | - | - | - |
|  | Tanzania | - | - | 2 | 3 | 2 | 2 | - | 2 | - | 3 | 2 | 2 | 1 | 2 | 2 | - | - |
|  | Uganda | 1 | 2 | 1 | - | 2 | 1 | - | 2 | 2 | 3 | - | 3 | 2 | 2 | 2 | - | - |
| Southern Africa | Botswana | - | 2 |  | - | - | - | - | 2 | - | - | - | 4 | - | 2 | - | - | 4 |
|  | Madagascar | 4 | 2 | 4 | - | 4 | 4 | - | 2 | - | - | 2 | 2 | - | 4 | - | - | - |
|  | Malawi | 1 | 2 | 1 | - | 2 | 2 | - | 2 | - | 3 | 2 | 2 | - | 3 | 3 | - | - |
|  | Mozambique | 2 | 3 | 2 | 3 | 2 | 2 | - | 2 | 3 | 3 | 3 | 2 | 4 | 4 | 3 | - | 4 |
|  | Zambia | - | 1 | 1 |  | 3 | 2 | - | 1 | - | - | - | 1 | - | - | 3 | - | 2 |
| Central Africa | Angola | 4 | 3 | 3 | - | - | - | - | 3 | - | - | 3 | 4 | - | - | - | - | - |
|  | Burundi | 2 | 2 | 2 | - | - | - | - | 2 | - | - | 2 | 2 | - | - | 2 | - | - |
|  | DRC Congo | 3 | 2 | 2 | - | - | - | - | 4 | - | - | 2 | 2 | - | - | 2 | - | - |
|  | Rwanda | 3 | 3 | 3 | - | - | - | - | 3 | - | - | 3 | 3 | - | - | - | - | - |
|  | Chad | - | - | 3 | - | 3 | 3 | - | 3 | 3 | - | - | 3 | 3 | 3 | - | - | 4 |
| West Africa | Benin | 1 | - | 1 | - | 1 | 1 | - | 1 | - | - | 1 | 1 |  |  | - | - | - |
|  | Burkina Faso | 2 | 2 | 2 | - | 2 | 3 | 4 | 2 | 2 | - | 3 | 2 | 2 | 2 | 2 | 3 | 4 |
|  | Cote d'Ivoir | - | - | 3 | - | 3 | - | - | 3 | - | - | - | 1 | - | - | - | - | - |
|  | Ghana | - | - | - | - | 1 | 1 | - | 1 | 1 | - | - | 1 | - | 1 | - | 1 | - |
|  | Guinea | - | 3 | 3 | - | 3 | - | - | 1 | - | - | 3 | 2 | - | - | - | - | - |
|  | Mali | 4 | 3 | 4 | 4 | 3 | 3 | 4 | 4 | 4 | 4 | 4 | 1 | 4 | 3 | 4 | 4 | 2 |
|  | Níger | - | - | - | - | 2 | 2 | - | 3 | 2 | - | 1 | 1 | 2 | 2 | - | - | - |
|  | Nigeria | - | - | - | - | 1 | 1 | - | 1 | 2 | - | - | 1 | 2 | 1 | - | - | 2 |
|  | Senegal | 3 | - | - | - | - | 4 | - | 3 | 2 | - | - | 1 |  | 2 | - | - | - |
|  | Togo | - | 4 | - | - | 3 | 4 | - | 3 | 4 | 4 | - | 4 | 4 | 4 | - | - | - |
