## Supplementary 4 for "Advancing Plant Health in Sub-Saharan Africa: Understanding National Agricultural Research Institutions capacities for Strategic Investment Priorities to Boost Genetic Gains Against Crop Biotic Stress"

**Supplementary table 4**: Proportion (%) of respondents engaged in plant health research activities on major crops across East, Southern, West, and Central Africa. The table highlights crop coverage among respondents, indicating varying levels of engagement by crop and region. Rice and maize were the most widely covered crops across regions, with 85% and 82% of respondents working on these crops, respectively. Moderate engagement was observed for beans, groundnut, cassava, banana, millet, and sorghum, with notable regional variation. Lower representation was recorded for crops such as chickpea, lentil, wheat, yam, and pigeonpea, reflecting more limited geographic and institutional focus. Overall, the data illustrates a strong emphasis on staple cereals and selected legumes, with uneven crop representation across regions.

| **Crop** | **Proportion (%) of respondents working on each crop per region** | | | |  |
| --- | --- | --- | --- | --- | --- |
|  | **East Africa** | **Southern Africa** | **West Africa** | **Central Africa** | **Across regions** |
| Rice | 67 | 86 | 88 | 100 | 85 |
| Maize | 67 | 86 | 76 | 100 | 82 |
| Beans | 33 | 86 | 35 | 75 | 57 |
| Groundnut | 67 | 57 | 71 | 25 | 55 |
| Cassava | 40 | 57 | 35 | 83 | 54 |
| Banana | 33 | 57 | 29 | 75 | 49 |
| Millet | 67 | 43 | 65 | 17 | 48 |
| Sorghum | 53 | 43 | 59 | 25 | 45 |
| Potato | 13 | 57 | 35 | 75 | 45 |
| Cowpea | 47 | 57 | 59 | 0 | 41 |
| Sweet potato | 47 | 43 | 24 | 42 | 39 |
| Sesame | 47 | 14 | 6 | 25 | 23 |
| Pigeonpea | 40 | 14 | 0 | 0 | 14 |
| Wheat | 13 | 29 | 0 | 0 | 10 |
| Chickpea | 27 | 0 | 0 | 0 | 7 |
| Yam | 0 | 0 | 18 | 0 | 4 |
| Lentil | 13 | 0 | 0 | 0 | 3 |
